# The smoking cessation drug cytisine requires systemically circulating estrogen for sex-specific neuroprotection in female parkinsonian mice

**DOI:** 10.1101/2024.03.21.586192

**Authors:** Sara M. Zarate, Roger C. Garcia, Gauri Pandey, Rahul Srinivasan

**Affiliations:** Department of Neuroscience & Experimental Therapeutics, Texas A&M University School of Medicine, 8447 Riverside Pkwy, Bryan, TX 77807-3260; Texas A&M Institute for Neuroscience (TAMIN)

**Keywords:** cytisine, smoking cessation, ovaries, estrogen, sex difference, aromatase

## Abstract

The smoking cessation drug cytisine exerts neuroprotection in substantia nigra pars compacta (SNc) dopaminergic (DA) neurons of female but not male 6-hydroxydopamine (6-OHDA) lesioned parkinsonian mice. To address the important question of whether circulating estrogen mediates this effect, we employ two mouse models aimed at depleting systemically circulating estrogen: (**i**) bilateral ovariectomy (OVX), and (**ii**) aromatase inhibition with systemically administered letrozole. In both models, depleting systemically circulating estrogen in female 6-OHDA lesioned parkinsonian mice results in the loss of cytisine-mediated neuroprotection as measured using apomorphine-induced contralateral rotations and SNc DA neurodegeneration. Our experiments also reveal that OVX alone exerts neuroprotection in SNc DA neurons due to compensatory changes not observed in the letrozole model, which underscores the importance of using independent models of estrogen depletion to study neuroprotection. Taken together, our findings suggest that the smoking cessation drug cytisine is a viable neuroprotective drug for pre-menopausal women with Parkinson’s disease.

## Introduction

Epidemiological studies over the past 60 years demonstrate that smokers and chronic tobacco users are at a 50% reduced risk for Parkinson’s disease (PD) ^1–4^. In this context, we and others have shown that at smoking relevant nanomolar concentrations, nicotine, which is the major addictive ingredient of tobacco, binds to and stabilizes the high sensitivity (α4)_2_(β2)_3_ stoichiometry of neuronal nicotinic acetylcholine receptors (nAChRs) within the endoplasmic reticulum (ER) of cells ^5–9^. In subsequent studies, we showed that nicotine chaperones nAChRs out of the ER via Sec24D-containing ER exit sites (ERES) into the cellular secretory pathway ^7,10,11^. Importantly, nicotine-mediated nAChR chaperoning in dopaminergic (DA) neurons is associated with an upregulation of Sec24D-containing ERES and an inhibition of all three signaling arms (XBP1, ATF6 and eIF2α) of the ER stress response pathway ^11^.

Given the established role of hyperactive ER stress in causing DA neuron loss during PD ^12,13^, our studies point to the general concept that nicotinic ligands can inhibit pathological ER stress signaling via nAChR chaperoning and ER exit site upregulation in DA neurons. Although this mechanism likely contributes to the neuroprotective effect of chronic tobacco use against PD, clinical trials employing nicotine as a disease-modifying treatment strategy for PD have failed, partly because of the adverse autonomic effects associated with large doses of systemic nicotine ^14–18^.

The smoking cessation drug cytisine is a partial agonist for α4β2 nAChRs with minimal side effects in humans ^19^. In addition, cytisine can bind to and chaperone α4β2 nAChRs at low nanomolar concentrations ^7,20,21^. Based on these favorable pharmacodynamic properties, we hypothesized that low nanomolar concentrations of cytisine could mediate neuroprotective effects against PD via the upregulation of nAChRs and ER exit sites and a consequent inhibition of the ER stress response in substantia nigra pars compacta (SNc) DA neurons. We tested this hypothesis *in vivo* by employing a preclinical mouse model of parkinsonism with unilateral injection of 6-hydroxydopamine (6-OHDA) into the dorsolateral striatum (DLS). Surprisingly, we found that when compared to saline treated controls, systemically administered low dose cytisine reduced the loss of SNc DA neurons only in female and not in male mice exposed to 6-OHDA ^21^. Furthermore, using mouse midbrain neuron-astrocyte co-cultures, we found that while cytisine inhibited two arms of ER stress signaling (ATF6 and XBP1), the active metabolite of estrogen,17-β-estradiol, inhibited the expression of CHOP which is a downstream apoptotic mediator of the third arm of ER stress signaling ^21^. Taken together, these findings suggest that estrogen acts in combination with cytisine to inhibit all three arms of ER stress response, thereby exerting sex-specific neuroprotection in female parkinsonian mice.

A major question that arises from our finding that cytisine exerts neuroprotection only in female parkinsonian mice is whether systemically circulating estrogen is essential for cytisine to exert neuroprotection of SNc DA neurons *in vivo*. This is a particularly important question because understanding the role of estrogen in cytisine-mediated neuroprotection could enable future use of cytisine as a neuroprotective treatment for both men and women with PD. Based on this rationale, the present study examines the extent to which systemically circulating estrogen is necessary for cytisine-mediated neuroprotection in a 6-OHDA-induced preclinical mouse model of parkinsonism. We employ two types of manipulations to deplete systemically circulating estrogen in female mice, *viz.* bilateral ovariectomy (OVX) and pharmacological inhibition of aromatase (AROM) with the AROM inhibitor, letrozole. In each of the two models with depleted systemically circulating estrogen, we show that cytisine fails to exert neuroprotection in female 6-OHDA exposed parkinsonian mice. Thus, our results suggest that systemically circulating estrogen is indeed essential for cytisine to exert neuroprotection in a unilateral 6-OHDA mouse model of parkinsonism. Our study has important implications for developing combined formulations of cytisine and estrogen analogs as a clinically translatable neuroprotective treatment for men and women with PD. Furthermore, this study underscores the importance of using multiple independent methods to assess the effects of circulating estrogen on neuroprotection in rodent models of PD.

## Materials and Methods

### Mice

Two- to three-month old female C57Bl6 mice were maintained on a 12 h light-dark cycle. Food and water were provided *ad libitum*. All experiments were performed in accordance with Texas A&M University IACUC regulations and protocols.

### Surgeries

#### Bilateral ovariectomy (OVX)

Mice were induced with 5% isoflurane and maintained with 2-3% isoflurane throughout the procedure. Fur was removed 2-3 cm lateral to the midline using clippers and the area was sterilized with 3 applications of iodine followed by 70% ethanol. Two 1-2 cm long incisions parallel to the midline were made through the skin and underlying muscle to access the ovaries. The uterine horns just proximal to the ovaries were tied off using polyglycolic acid absorbable synthetic sutures. The ovaries and surrounding fat were then excised. For sham surgeries, the ovaries were accessed but not removed. Post-operative monitoring was performed for one week after OVX to ensure proper wound healing. Only mice that showed no signs of pain or distress for one week following OVX were used in experiments.

#### Unilateral lesion of the dorsolateral striatum (DLS) with 6-OHDA

All mice were anesthetized with isoflurane as described for OVX, and unilateral stereotaxic injection of 10 ug 6-OHDA in the dorsolateral striatum was performed as previously described ^21^. Briefly, 2 µl of a 5 mg/ml stock solution of 6-OHDA in 0.9% saline with 0.2% ascorbic acid was injected into the DLS at a flow rate of 1 μl/min. Coordinates for the injection site were 0.8 mm anterior to bregma, 2.0 mm lateral to the midline, and 2.4 mm ventral to the pial surface. In the week following 6-OHDA lesion, all mice were monitored daily for excessive weight loss prior to resuming behavioral testing.

#### Apomorphine induced rotational behavior

Apomorphine induced rotations were performed as previously described ^21^. Mice were habituated in separate 5-gallon buckets for 15 minutes and then received i.p. injection of 0.5 mg/kg apomorphine (Sigma, cat# 1041008) prepared in 0.9% saline. Mice were recorded for 15 minutes to capture the total number of contralateral rotations. Contralateral rotations were analyzed using Ethovision XT (Noldus, Leesburg, VA) and arena sizes, trial settings, and detection settings, were maintained constant for all mice across all weeks of behavior.

#### Methods to deplete estrogen in mice

Systemic estrogen was depleted using two independent methods: (**i**) bilateral OVX (Experiment 1) and (**ii**) inhibition of aromatase with letrozole (Experiment 2). For both experiments, mice were injected intraperitoneally (i.p.) with either saline (control) or cytisine and were assayed for apomorphine-induced contralateral rotations before and after 6-OHDA lesion. Sections below describe detailed protocols for the two experiments to deplete estrogen.

#### Experiment 1

C57Bl6 female mice (2-3 months) received either an OVX or sham surgery. As a treatment, all mice received alternate day intraperitoneal (i.p.) injections of either 200 µl 0.9% saline or 0.2 mg/kg cytisine in 0.9% saline for two weeks after OVX and throughout the remainder of the experiment. The mice were randomly separated into the following treatment groups: (**i**) sham operated females treated with saline (Sal SO), (**ii**) sham operated females treated with cytisine (Cyt SO), (**iii**) OVX females treated with saline (Sal OVX), or (**iv**) OVX females treated with cytisine (Cyt OVX). Behavior assays were performed prior to OVX, prior to 6-OHDA lesion, and two weeks post 6-OHDA lesion.

#### Experiment 2

C57Bl6 female mice (2-3 months) were sorted into the following treatment groups: (**i**) saline treated, (**ii**) letrozole treated (Let), or (**iii**) cytisine and letrozole treated (Cyt + Let). One week prior to 6-OHDA lesion, the saline and letrozole group received alternate day i.p. injections of 100 µl of 0.9% saline, while Cyt + Let group received 0.2 mg/kg cytisine diluted in 0.9% saline. After 6-OHDA lesion, saline mice received dual injections of 100 µl 0.9% saline and 100 µl of corn oil + 1% DMSO. Letrozole mice received 100 µl 0.9% saline and 100 µl of 1 mg/kg letrozole diluted in corn oil + 1% DMSO. Cyt + Let mice received 1 mg/kg letrozole and 0.2 mg/kg cytisine. Apomorphine rotation behavior assays were performed before and after 6-OHDA lesion.

### Histology

#### Tissue Collection

After the final behavior assays were performed, all mice were sacrificed with isoflurane anesthesia and transcardially perfused with 1X PBS followed by 10% formalin. Following perfusion, the uterus and brain were extracted from each animal. Uterine weights were measured using precision balances.

Extracted brains were placed in 10% formalin overnight at 4°C followed by 30% sucrose until the time of sectioning. Using a Leica CM1850 cryostat (Leica, Deer Park, IL), 40 µm coronal sections containing the midbrain were obtained. All sections were collected in 0.01% sodium azide (Sigma, cat# S2002) and stored sequentially in 96 well plates to maintain the exact rostrocaudal positioning of the midbrain sections.

#### Quantification of SNc DA neuron loss

The extent of 6-OHDA induced SNc DA neurodegeneration was assessed by obtaining ratios of lesioned and unlesioned SNc tyrosine hydroxylase (TH) fluorescent intensities from mouse midbrain sections as previously described ^21^. Briefly, 10 midbrain sections were selected at 120 µm intervals from each mouse for TH immunostaining. Following a wash in PBS, sections were blocked and permeabilized in 10% normal goat serum (Abcam, ab7481) and 0.5% Triton X-100 rocking for 45 min at RT. Sections were washed twice with PBS before incubation overnight at 4°C with chicken anti-TH (1:2000, Abcam cat# ab76442) in 1% NGS + 0.05% Triton X-100. The next day sections were washed twice with PBS followed by incubation in goat anti-chicken AlexaFluor 594 (1:2000, Abcam, ab150176) for 1 h at RT.

For each midbrain section, rectangular 10 mm^2^ regions of interest (ROIs) were drawn around the midbrain such that the lesioned and unlesioned SNc could be obtained in a single image. Each image contained a 20 μm thick optical z-stack and was aquired using an Olympus VS120 equipped with a 10x objective. Of the 10 immunostained midbrain sections, only 8 sections whereby the SNc and VTA could be visually separated were selected for imaging. Imaging parameters were maintained across all sections and mice. Z-stacks were then sum projected using Fiji software and individual ROIs were manually drawn around the lesioned and unlesioned SNc with the polygon tool. Each ROI was thresholded to select TH^+^ cell bodies and fibers to obtain integrated fluorescent intensities for each ROI. Fluorescent intensities across 8 midbrain sections per mouse for either the lesioned or unlesioned SNc were totaled and ratios of lesioned to unlesioned SNc fluorescent intensities were compared.

#### Quantification of glial fibrillary acid protein (GFAP) and aromatase (AROM) staining

To assess the effects of estrogen depletion on astrocyte reactivity and brain derived estrogen, we respectively quantified GFAP and AROM expression in mouse midbrains. For GFAP and AROM quantification, 4 midbrain sections from mice in Experiment 1were selected at 440 µm intervals between sections. Sections were washed in 1X PBS, blocked and permeabilized in 10% normal goat serum (Abcam, ab7481) and 0.5% Triton X-100 with rocking for 45 min at RT. Sections were washed twice with PBS before incubation overnight at 4°C with rabbit anti-aromatase (1:2000, Abcam cat # ab18995), chicken anti-TH (1:2000, Abcam cat# ab76442), and mouse anti-GFAP (1:2000, Invitrogen cat# 14-9892-82) in 1% NGS + 0.05% Triton X-100. The next day, sections were washed twice with PBS followed by incubation in goat anti-rabbit AlexaFluor 488 (1:2000, Abcam cat# ab150077), goat anti-chicken AlexaFluor 594 (1:2000, Abcam cat# ab150176), and goat anti-mouse AlexaFluor 647 (1:2000, Abcam cat# ab150115) for 1 h at RT.

Two z-stacks from the lesioned and unlesioned SNc were acquired from each section using an Olympus FV3000 confocal microscope equipped with LED excitation at 488, 594 and 647 nm and a 60x oil objective lens. Each z-stack was composed of 25 optical sections with a step size of 0.49 µm. All imaging parameters were maintained across treatment groups and imaging days. Using Fiji, a max projection was made from each z-stack and a histogram range that subtracted background fluorescence was applied to each image. The same histogram range was used for all images. Images were then thresholded to select all the pixels in the image and the threshold was applied to the image to create a binary mask. The binary mask was then combined with the original image using the AND image calculator function in Fiji. The AND function results in an image containing only the pixels present in both images. SNc fluorescent intensities for GFAP and AROM from AND images were obtained for each treatment group.

#### Statistics

Origin 2022 (OriginLab, Northampton, MA) was used for statistical analysis. All data sets were first tested for normality. For comparisons between 2 treatment groups, normally distributed data were analyzed with two-sample t-tests. A two-way mixed design ANOVA was performed to determine the between subject effects of a fixed factor (treatment group) on body weight with a repeated measure designated as average body weight for a given week ^22^. A one-way ANOVA test was performed to compare three or more treatment groups after 6-OHDA lesion ^23^. Statistical significance was considered as *p* < 0.05. The types of statistical tests used are detailed in results, and sampling numbers for each experiment are detailed in each figure legend.

## Results

### Cytisine fails to reduce apomorphine-induced contralateral rotations in 6-OHDA lesioned mice with bilateral OVX

To determine if the depletion of systemically circulating estrogen could prevent neuroprotection by cytisine in 6-OHDA lesioned parkinsonian mice, bilateral OVX was performed in 2 – 3-month old female mice. Sham operated (SO) mice with intact ovaries were used as controls (Figure 1A). Since OVX-induced depletion of circulating estrogen is known to cause an increase in body weight, along with a decrease in uterine weight ^24^, we compared whole body and uterine weights between control SO and OVX mice. Compared to SO mice, OVX mice showed a significant 12% increase in body weight at two weeks post-OVX surgery, which persisted throughout the remainder of the experiment (F_(3,65)_ = 6.61, *p* = 0.01; two-way mixed design ANOVA) (Figure 1B). Additionally, 5 weeks post-OVX, uterine weights were significantly reduced by ∼80% in OVX mice when compared to SO mice (F_(3,61)_ = 14.6, *p* < 0.0001; one-way ANOVA) (Figure 1C). These data indicate successful depletion of circulating estrogen in bilateral OVX mice.

**Figure 1.**
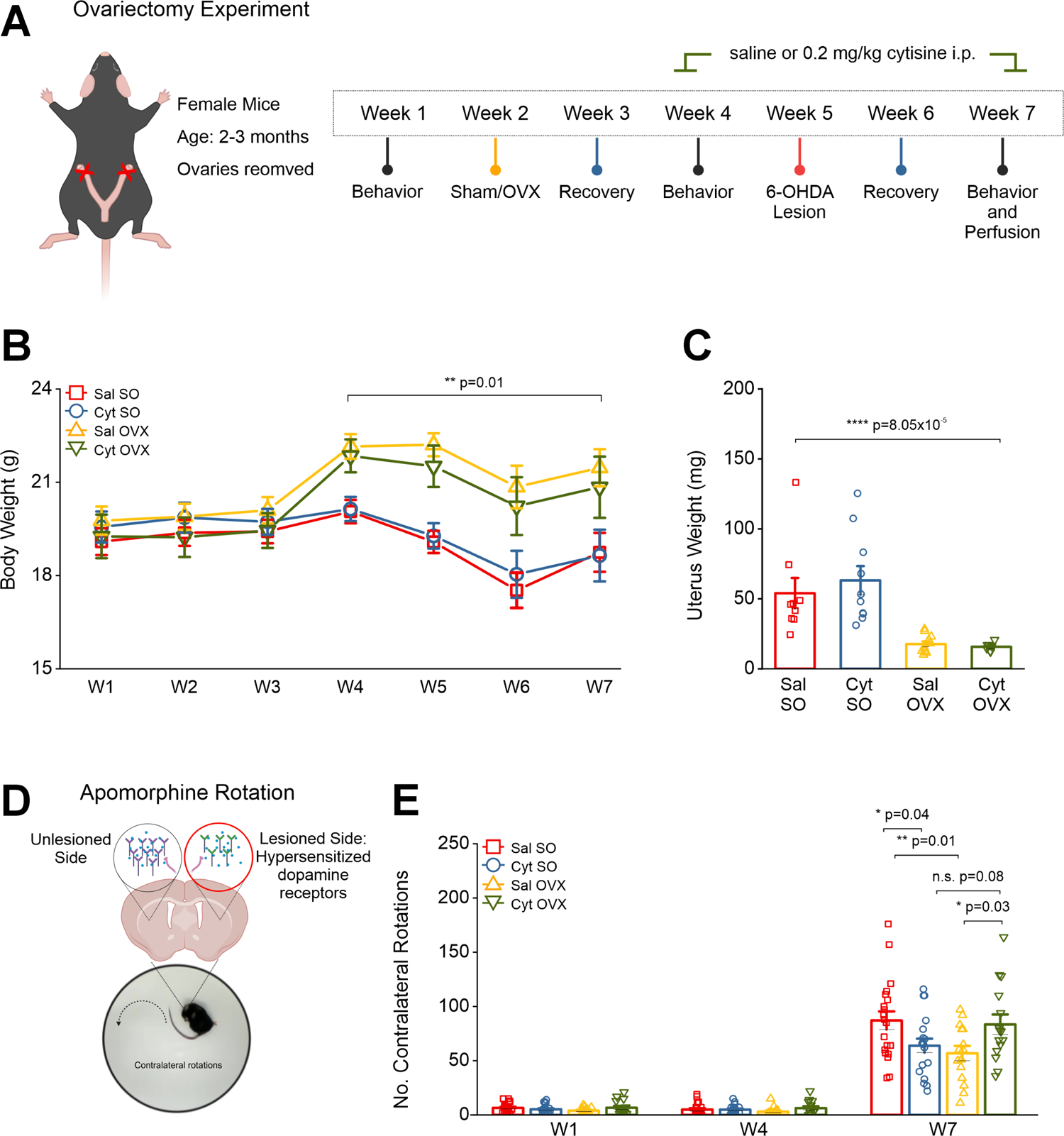
OVX induces long lasting changes to body and uterine weight. (A) Schematic showing the protocol and timeline for OVX experiment in female mice (B) Comparison of body weight changes across the experiment timeline for each treatment group (C) Comparison of uterine weights collected at the end of the experiment, 5 weeks after OVX (D) Schematic showing the rationale for apormophine-induced contralateral rotations that occurs due to hypersensitized dopamine receptors in the 6-OHDA lesioned DLS (E) Bar graph showing the number of contralateral rotations counted in 15 min after i.p. injection with 0.5 mg/kg apomorphine for weeks 1, 4, and 7. Error bars are S.E.M. For panel B the p value was derived from a two-way mixed design ANOVA considering sham/OVX by time. For panel C, p values were derived from a one-way ANOVA test for all treatment groups. Comparison for increased rotations in W1-7 p value is derived from a one-way repeated measures ANOVA. For between group comparisons on W7 p values are based on two-sample t-tests n = 20 saline SO, n = 19 cytisine SO, n = 15 saline OVX, and n = 15 cytisine OVX female mice.

As shown in Figure 1A, SO and OVX mice received alternate day i.p. injections of either 0.9% saline (Sal) or 0.2 mg/kg cytisine (Cyt) starting in week 4, followed by a unilateral striatal lesion with stereotaxic injection of 6-OHDA into the dorsolateral striatum in week 5. Following 6-OHDA striatal lesions, saline treated sham operated (Sal SO) mice showed a 12-fold increase in apomorphine rotations when compared to baseline from 6.5 ± 0.98 rotations in W1 to 87.1 ± 8.35 rotations in W7 (F_(1.03,19.62)_ = 103.22; *p* = 2.4 x 10^-9^; one-way repeated measures ANOVA) (Figure 1E). Consistent with our prior findings ^21^, SO mice treated with cytisine (Cyt SO) showed significantly reduced apomorphine rotations when compared to control Sal SO mice (Cyt SO: 63.84 ± 6.46 rotations; Sal SO: 87.1 ± 8.35 rotations, *p* = 0.04; two-sample t-test) (Figure 1E). By contrast, when compared to Sal OVX mice, Cyt OVX mice demonstrated a significant increase in rotations after 6-OHDA lesion (Sal OVX: 56.8 ± 6.8 rotations; Cyt OVX: 83.4 ± 9.22 rotations, *p* = 0.08; two-sample t-test) (Figure 1E). These data indicate that cytisine fails to reduce unilateral apomorphine rotational behavior in mice with depletion of circulating estrogen. It should be noted that Sal OVX mice demonstrated significantly reduced apomorphine rotations when compared to Sal SO mice (Sal SO: 87.1 ± 8.35 rotations; Sal OVX: 56.8 ± 6.8 rotations, *p* = 0.01; two-sample t-test) (Figure 1E), which suggests that OVX alone reduces apomorphine-induced contralateral rotational behavior in mice with unilateral 6-OHDA lesions in the striatum.

### Cytisine does not prevent 6-OHDA-induced loss of SNc DA neurons in OVX mice

Having observed significant differences in contralateral apomorphine rotations among experimental groups (Figure 1E), we sought to determine the extent to which cytisine and/or OVX treatment affects 6-OHDA-induced loss of SNc DA neurons. To do this, all mice were perfused in week 7 of the experimental protocol shown in Figure 1A. Forty-micron midbrain sections were obtained along the rostro-caudal axis from these mice and immunostained for TH (Figure 2A and 2B). To quantify the extent of midbrain SNc DA neuron loss for each of these mice, we obtained integrated TH intensities for the lesioned and unlesioned SNc (Figure 2C) and lesioned / unlesioned SNc TH integrated intensity ratios (Figure 2D) for all mice in each experimental group as described in methods. We found that the ratio of lesioned to unlesioned SNc TH intensity was significantly higher in Cyt SO mice when compared to Sal SO (Sal SO, TH ratio: 0.06 ± 0.012 and Cyt SO, TH ratio: 0.15 ± 0.031, *p* = 0.03; two-sample t-test) (Figure 2D). These data confirm our previous finding that cytisine alone exerts a neuroprotective effect on 6-OHDA lesioned SNc DA neurons in female mice with intact ovaries ^21^.

**Figure 2.**
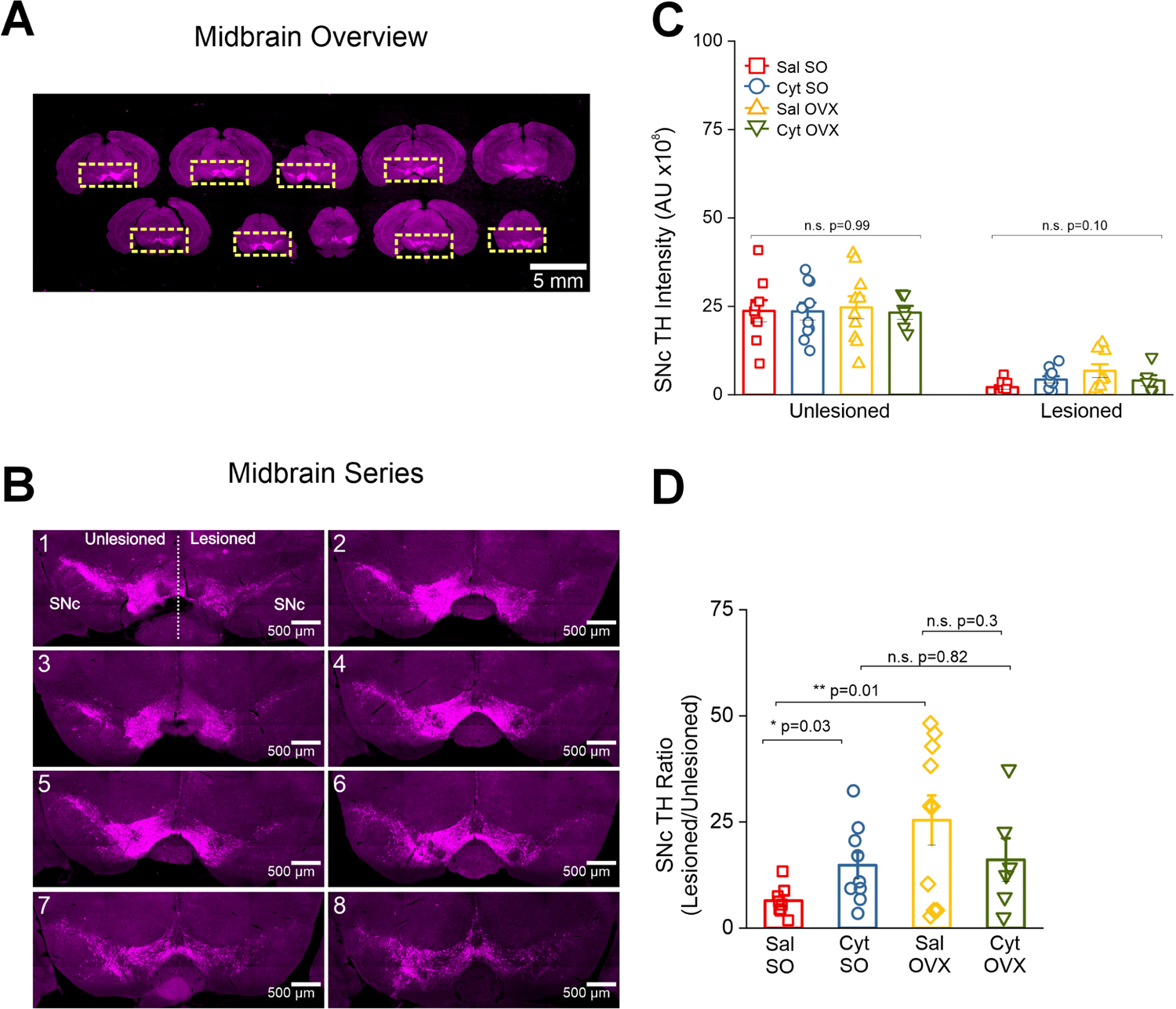
OVX does not reverse cytisine induced neuroprotection after 6-OHDA lesion. (A) Low magnification representative images of serial midbrain sections at 120 μm intervals. Only sections where the boundary between the SNc and VTA was clearly defined were selected for analysis. ROIs were manually demarcated such that the unlesioned and lesioned midbrain can be viewed in a single image. Scale bar = 5 mm (B) Representative images of a midbrain series from a single mouse containing 8 TH labeled midbrain sections used for analysis. Scale bar = 500 µm (C) Bar graphs comparing total TH intensity in either the unlesioned or lesioned SNc for all treatment groups. Each data point is the sum of TH intensity measured for either the unlesioned or lesioned SNc in 8 images per mouse (D) Bar graphs comparing TH intensity ratios for all treatment groups. Each data point is the total lesioned SNc TH intensity by the total unlesioned SNc TH intensity. Error bars are S.E.M. For panel C, the p values for the unlesioned and lesioned comparisons are derived from a one-way ANOVA comparing each treatment group. For panel D, p values are derived from two-sample t-tests. n = 9 saline SO, n = 10 cytisine SO, n = 10 saline OVX, and n = 6 cytisine OVX female mice.

By contrast, when compared to Sal OVX mice, Cyt OVX mice failed to show a significant increase in SNc TH ratios (Sal OVX, TH ratio: 0.24 ± 0.05 and Cyt OVX, TH ratio: 0.16 ± 0.05, *p* = 0.3; two-sample t-test) (Figure 2D). Taken together, these data show that OVX-induced depletion of circulating estrogen prevents the neuroprotective effect of cytisine in midbrain SNc DA neurons. It should be noted that when compared to Sal SO mice, Sal OVX mice showed a significant 2.7-fold increase in TH intensity ratio after 6-OHDA lesion (Sal SO, TH ratio: 0.06 ± 0.012 and Sal OVX, TH ratio: 0.24 ± 0.05, *p* = 0.01; two-sample t-test) (Figure 2D), which suggests a neuroprotective effect of OVX alone.

### Cytisine does not decrease astrocyte reactivity in the 6-OHDA lesioned SNc of OVX mice

Increased astrocyte reactivity occurs as a consequence of ongoing DA neurodegeneration within the SNc, which can accelerate the neurodegenerative process in PD ^25–27^. Based on this rationale, we sought to quantify changes in GFAP intensity within the 6-OHDA-lesioned SNc of mice from all experimental groups *viz.* Sal SO, Cyt SO, Sal OVX and Cyt OVX. Midbrain sections were stained for GFAP and confocal images of the lesioned SNc were obtained for quantification of GFAP intensity in each mouse (Figure 3A). Compared to Sal SO mice, Cyt SO mice showed a significant decrease in GFAP intensity within the lesioned SNc (Sal SO, GFAP intensity: 14.90 ± 1.16 and Cyt SO, GFAP Intensity: 7.36 ± 1.65, *p* = 0.03; one-way ANOVA with post-hoc Tukey test). (Figure 3B), which correlates with the neuroprotective effect of cytisine shown in Figure 2D. However, when compared to Sal SO mice, Sal OVX mice showed no significant difference in GFAP intensity (Sal SO, GFAP intensity:14.90 ± 1.16 and Sal OVX, GFAP intensity: 10.22 ± 2.27, *p* = 0.23; one-way ANOVA with post-hoc Tukey test) (Figure 3B). Importantly, when compared to Sal OVX mice, Cyt OVX mice did not show a further decrease in GFAP intensity within the lesioned SNc (Sal OVX, GFAP intensity: 10.22 ± 2.27 and Cyt OVX, GFAP intensity: 7.35 ± 1.34, *p* = 0.62; one-way ANOVA with post-hoc Tukey test) (Figure 3B). This finding provides additional evidence that the neuroprotective effect of cytisine in the SNc requires circulating estrogen.

**Figure 3.**
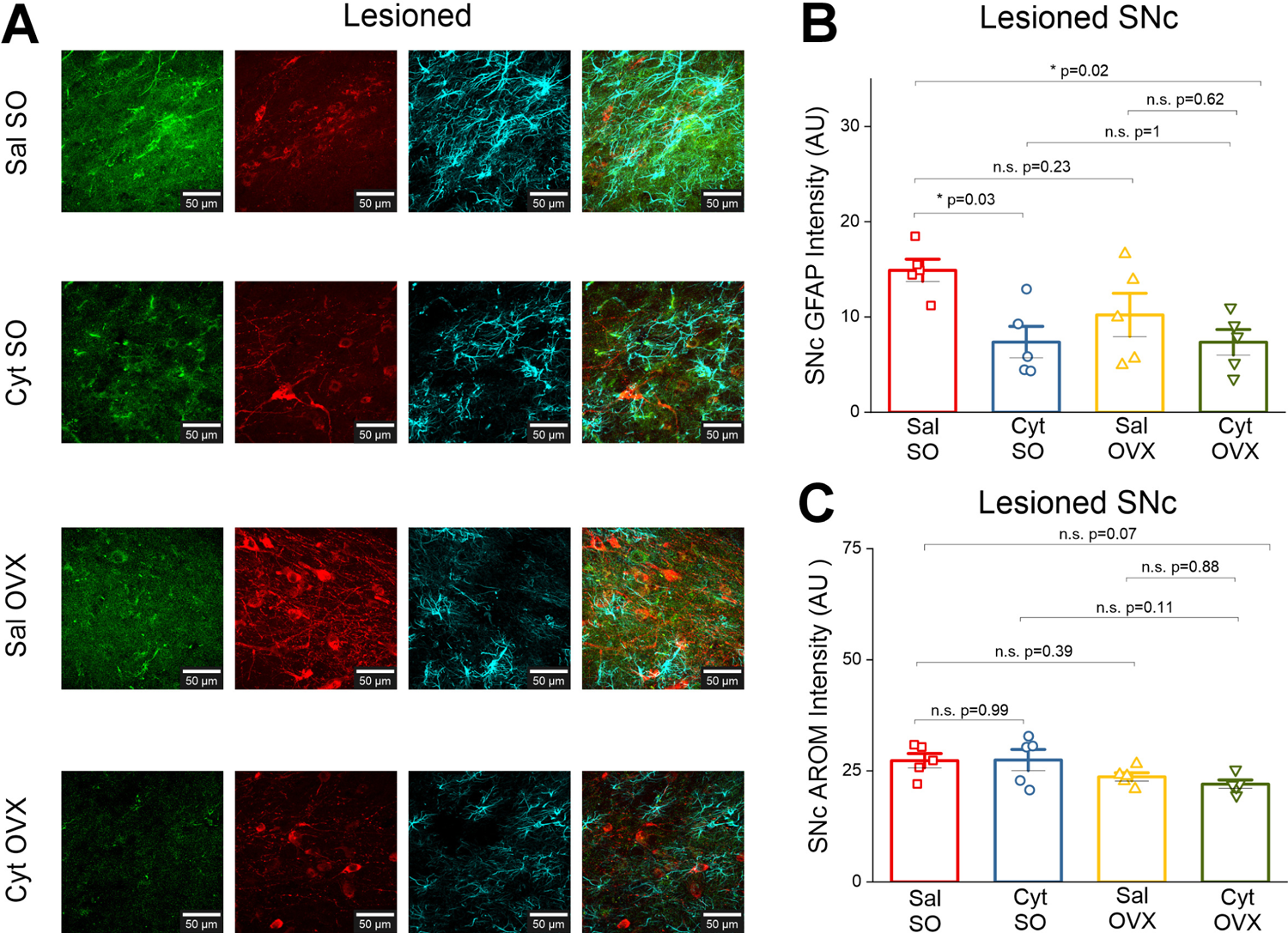
Cytisine neuroprotection in OVX mice is independent of glial derived estrogen. (A) Representative confocal images from the lesioned SNc across treatment groups. All sections were stained for AROM (green), TH (red), and GFAP (cyan). Scale bar = 25 µm (B) Bar graph comparing total GFAP intensity in the lesioned SNc for all treatment groups (C) Bar graph comparing total AROM intensity in the lesioned SNc for all treatment groups. Each data point is the average of either GFAP or AROM intensity measured from 4 images for the unlesioned SNc and 4 images for the lesioned SNc. Error bars are S.E.M. P values are derived from a one-way ANOVA and Tukey post hoc comparisons between treatment groups. n = 5 saline SO mice, n = 5 cytisine SO mice, n = 5 saline OVX mice, and n = 5 cytisine OVX mice.

### Brain-derived estrogen does not contribute to cytisine-mediated neuroprotection in the 6-OHDA lesioned SNc

Results thus far show that depletion of circulating estrogen via OVX results in a loss of neuroprotection by cytisine. Cells in the brain are steroidogenic and express aromatase (AROM), which is the primary cytochrome P450 enzyme (CYP19A1) that synthesizes 17-β-estradiol in the brain ^28^.

Therefore, to determine the contribution of brain-derived estrogen to cytisine-mediated neuroprotection we sought to quantify changes in AROM expression in the lesioned SNc of mice from all experimental groups *viz.* Sal SO, Cyt SO, Sal OVX and Cyt OVX. Midbrain sections from mice in each experimental group were stained for AROM (Figure 3A) and AROM intensity was quantified from confocal images of the lesioned SNc in these mice as described in methods. Midbrain AROM intensity analysis revealed that neither cytisine treatment nor OVX resulted in significant changes in aromatase expression (Sal SO, AROM intensity: 27.29 ± 1.62; Cyt SO, AROM intensity: 27.44 ± 2.38; Sal OVX, AROM intensity: 23.66 ± 0.92, Cyt OVX, AROM intensity: 22.02 ± 0.93, *p* = 0.07; one-way ANOVA) (Figure 3C). These data suggest that the loss of cytisine-mediated neuroprotection in OVX mice stems from a depletion of circulating and not brain-derived estrogen.

### Depletion of estrogen with the AROM inhibitor letrozole prevents cytisine-induced neuroprotection of SNc DA neurons

We have shown that the depletion of systemically circulating estrogen via OVX prevents cytisine-mediated neuroprotection of SNc DA neurons (Figures 1 – 3). However, when compared to mice with intact ovaries, we find that OVX alone exerts neuroprotection in the SNc DA neurons of 6-OHDA lesioned mice (Figure 2D). Based on these findings, we sought to assess cytisine neuroprotection using systemic AROM inhibition as a model of estrogen depletion, which would reduce the long-term compensatory effects that are seen with OVX.

We administered 1 mg/kg of the AROM inhibitor, letrozole (Let) i.p to mice for two weeks on alternate days, starting one day after striatal 6-OHDA lesion (Figure 4A). The three experimental groups of 6-OHDA lesioned mice were as follows: (**i**) control 6-OHDA lesioned mice with only i.p. saline, (**ii**) control 6-OHDA lesioned mice with only 1 mg/kg Let i.p., and (**iii**) experimental 6-OHDA lesioned mice with 1 mg/kg of Let and 0.2 mg/kg Cyt i.p. We found that when compared to saline treated mice, Let treatment did not result in a significant difference in body or uterus weight (*p* = 0.92, one way ANOVA) (Figures 4B and C), suggesting minimal long-term compensatory changes following Let treatment. After 6-OHDA lesions, all three groups of mice were compared for apomorphine-induced rotations in week 4 (Figure 4D). We found no significant difference in the number of contralateral rotations between the three groups of mice (saline vs let vs cyt + let: *p* = 0.92, one way ANOVA) (Figure 4D). We then stained midbrain sections from all mice to determine DA neuron survival after 6-OHDA lesion (Figure 4E). No difference was detected when TH intensity ratios were compared across treatment groups (F_(2,20)_ = 1.27, *p* = 0.3, one-way ANOVA) (Figures 4F and G). Together, these data demonstrate that the acute depletion of estrogen in female mice with intact ovaries also prevents cytisine induced neuroprotection of 6-OHDA lesioned SNc DA neurons.

**Figure 4.**
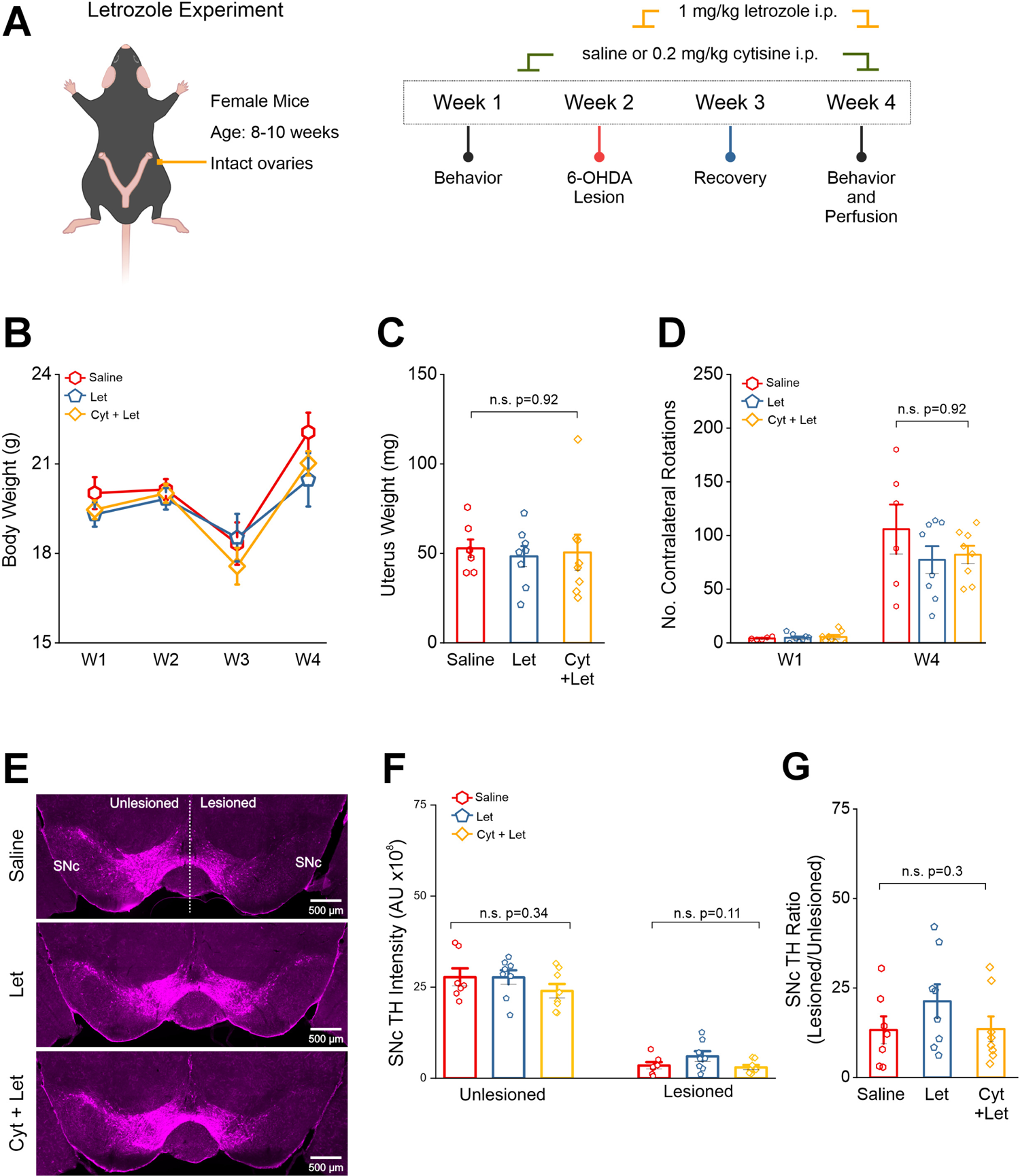
Cytisine neuroprotection is reversed by aromatase inhibition. (A) Schematic of protocol for behavior assays (B) Comparison of body weight changes across the experiment timeline for each treatment group (C) Comparison of uterine weights collected at the end of the experiment (D) Bar graph showing the number of contralateral rotations counted in 15 min after i.p. injection with 0.5 mg/kg apomorphine for weeks 1 and 4 (E) Representative images of a single midbrain section from each treatment group. Scale bar = 500 µm (F) Bar graphs comparing total TH intensity in either the unlesioned or lesioned SNc for all treatment groups. Each data point is the sum of TH intensity measured for either the unlesioned or lesioned SNc in 8 images per mouse (G) Bar graphs comparing TH intensity ratios for all treatment groups. Error bars are S.E.M. For panels C, D, and G, p values were derived from a one-way ANOVA test for all treatment groups. For panel F, the p value is derived from a one-way ANOVA comparing each treatment group in the unlesioned or lesioned SNc. n = 7 saline mice, n = 8 letrozole mice (let), and n = 8 cytisine + letrozole mice (cyt+let).

## Discussion

In this study, we determine the extent to which the smoking cessation drug cytisine requires systemically circulating estrogen for neuroprotection against 6-OHDA-induced SNc DA neuron loss in female mice. To do this, we employ two different mouse models that deplete circulating estrogen in 6-OHDA lesioned female mice: (**i**) bilateral OVX, and (**ii**) acute pharmacological inhibition of AROM with letrozole. In each of these models, we show that when compared to estrogen depleted mice, pre-treatment with cytisine in a background of systemic estrogen depletion results in the loss of cytisine-mediated neuroprotection in SNc DA neurons (Figures 2D and 4G). Importantly, the current study recapitulates our previous finding that cytisine exerts neuroprotection in female mice with intact ovaries (Figure 2D) ^21^.

The use of two types of models to deplete systemically circulating estrogen is an important aspect of this study because this approach addresses the possibility of model-specific idiosyncrasies that can lead to incorrect interpretation of data. One such idiosyncrasy is that when compared to sham operated mice with intact ovaries, OVX mice with unilateral striatal 6-OHDA lesions display significantly reduced contralateral rotations with apomorphine, which is indicative of neuroprotection (Figure 1E). This effect could partly be explained by OVX-mediated compensatory changes in the striatum, specifically a decrease in the expression of striatal dopamine type 1 (D1) receptors ^29,30^. Apomorphine rotations are induced by activation of hypersensitive dopamine receptors in striatal medium spiny neurons (Figure 1D), with the number of rotations providing an indirect measure of SNc DA neuron loss after unilateral striatal lesion ^31,32^. Thus, an OVX-induced compensatory decrease in striatal D1 receptors likely contributes to reducing the total number of contralateral rotations with apomorphine. Extending these data, we find that pre-treatment of OVX mice with cytisine causes a significant increase in apomorphine rotations when compared to mice with OVX alone (Figure 1E). Our interpretation of this result is that chronic cytisine treatment upregulates α4β2* nAChRs (* indicates additional uncharacterized nAChR subunits in the ion channel pentamer) in SNc DA terminals within the striatum, which would in turn increase the basal release of dopamine in the striatum ^33^. Therefore, it follows that the activation of striatal D1 receptors by a cytisine-induced increase in basal dopamine release on the lesioned side would generally increase apomorphine-induced contralateral rotations in cytisine-treated OVX mice with unilateral loss of SNc DA neurons.

In addition to reducing apomorphine rotations, we show that OVX alone exerts a neuroprotective effect in 6-OHDA-lesioned SNc DA neurons of mice (Figure 2D). This observation can be explained by our finding that OVX causes a 50% reduction in SNc astrocyte reactivity, measured using SNc GFAP expression as a readout (Figure 3B). Our finding that OVX-induced depletion of estrogen can reduce GFAP expression in SNc astrocytes is supported by a previous study showing that OVX in mice robustly alters GFAP promoter activity in the central nervous system following ischemia ^34^, suggesting that systemically circulating estrogen can indeed exert an effect on astrocyte reactivity during brain injury.

Furthermore, the idea that an OVX-induced decrease in astrocyte reactivity can be specifically neuroprotective in the SNc is supported by our prior demonstration that astrocyte processes in the SNc completely envelop DA neuron somata ^35^, which places SNc astrocytes in a unique morphological position to exert direct effects on SNc DA neuron survival. In this context, one future question that remains to be explored is the extent to which cytisine can directly alter SNc astrocyte function during the process of neurodegeneration. For example, cytisine could cause astrocytes to increase their secretion of antioxidants ^36^, mitigate mitochondrial damage in SNc DA neurons ^37^, or increase phagocytosis of abnormal protein aggregates ^38^.

It should be noted that OVX results in a 70 to 80% loss of TH content in the SNc after 6-OHDA lesion, which is not attenuated by the pretreatment of OVX mice with cytisine (Figure 2D). This suggests that cytisine requires systemically circulating estrogen for neuroprotection. However, it is also possible that 6-OHDA lesions along with OVX and/or cytisine can upregulate brain-derived AROM, thereby causing neuroprotection by increasing local synthesis of 17-β-estradiol within the SNc. We rule out this possibility by showing that OVX alone does not alter AROM expression in the SNc when compared to mice with intact ovaries (Figure 3C). Furthermore, cytisine does not significantly alter expression of AROM in SO or OVX groups. Based on these data, we infer that when compared to OVX only mice, the observed loss of cytisine-mediated SNc neuroprotection by cytisine in the background of OVX is due to depletion of systemically circulating estrogen rather than changes in production of brain-derived estrogen.

The fact that OVX results in a neuroprotective effect in the lesioned SNc motivated us to employ a second model for estrogen depletion using letrozole, which is an AROM inhibitor. The absence of an abnormal increase in body weight or a decrease in uterine weight (Figures 4B and 4C), as well as a lack of changes in apomorphine rotations (Figure 4D) in letrozole treated mice supports our rationale that the strategy of acutely depleting estrogen immediately after 6-OHDA lesions mitigates long-term compensatory effects in mice. Importantly, in a background of letrozole-induced acute estrogen depletion, we found that cytisine fails to exert neuroprotective effects in SNc DA neurons (Figure 4D – G). Taken together, based on our findings that cytisine exerts neuroprotection in mice with intact ovaries, but does not exert additional neuroprotection in OVX or letrozole treated mice, we conclude that cytisine requires systemically circulating estrogen for exerting neuroprotection in 6-OHDA lesioned female mice.

One limitation of this study is that we do not delineate downstream mechanisms by which circulating estrogen can act in combination with cytisine to prevent the loss of SNc DA neurons. In light of our previous study, it is likely that estrogen exerts neuroprotection by inhibiting expression of the pro-apoptotic ER stress molecule, CHOP ^21^. However, other downstream neuroprotective mechanisms such as ERES upregulation via estrogen-mediated chaperoning of nAChRs or direct genetic / epigenetic effects of estrogen in SNc DA neurons cannot be ruled out as potential neuroprotective mechanisms for the combined neuroprotective effects of estrogen and cytisine in female mice. Regardless of downstream mechanism(s) involved, the current study bolsters our view that the smoking cessation drug cytisine prevents SNc DA neurons loss by acting in combination with circulating estrogen, and can therefore be regarded as a potential neuroprotective drug for pre-menopausal women with PD. In future studies, we will assess the extent to which a combination of cytisine along with systemically administered estrogen analogues can exert neuroprotection in SNc DA neurons of male parkinsonian mice.

## Author contributions

SMZ performed experiments, analyzed data, interpreted results and wrote the first draft of the manuscript. RCG and GP performed experiments and analyzed data. RS conceptualized the study, designed experiments, supervised SMZ, RCG and GP, provided resources and funding for the study, and wrote and edited the manuscript.

## Acknowledgements

This study was funded by a National Institutes of Health (NIH) research grant, R01NS115809 to RS.

## Competing interests

All authors declare no financial or non-financial competing interests.

## Data Availability

The datasets used and/or analyzed during the current study are available from the corresponding author on reasonable request.

## Notes

### Competing Interest Statement

The authors have declared no competing interest.

